# Aging-associated augmentation of gut microbiome virulence capability drives sepsis severity

**DOI:** 10.1101/2023.01.10.523523

**Authors:** James F. Colbert, Joshua M. Kirsch, Christopher L. Erzen, Christophe J. Langouët-Astrié, Grace E. Thompson, Sarah A. McMurtry, Jennifer M. Kofonow, Charles E. Robertson, Elizabeth J. Kovacs, Ryan C. Sullivan, Joseph A. Hippensteel, Namrata V. Sawant, Nicole J. De Nisco, Bruce D. McCollister, Robert S. Schwartz, Alexander R. Horswill, Daniel N. Frank, Breck A. Duerkop, Eric P. Schmidt

## Abstract

Prior research has focused on host factors as mediators of exaggerated sepsis-associated morbidity and mortality in older adults. This focus on the host, however, has failed to identify therapies that improve sepsis outcomes in the elderly. We hypothesized that the increased susceptibility of the aging population to sepsis is not only a function of the host, but also reflects longevity-associated changes in the virulence of gut pathobionts. We utilized two complementary models of gut microbiota-induced experimental sepsis to establish the aged gut microbiome as a key pathophysiologic driver of heightened disease severity. Further murine and human investigations into these polymicrobial bacterial communities demonstrated that age was associated with only subtle shifts in ecological composition, but an overabundance of genomic virulence factors that have functional consequence on host immune evasion.

**One Sentence Summary:** The severity of sepsis in the aged host is in part mediated by longevity-associated increases in gut microbial virulence.

## INTRODUCTION

Sepsis is a common, lethal, and incompletely understood clinical entity that disproportionally impacts the aging population ^1,2^. Multiple epidemiologic studies have demonstrated a strong association of longevity with both sepsis incidence and case fatality rate ^3,4^. Older patients are particularly likely to develop sepsis caused by bacteria originating in the gut microbiota, via clinical syndromes such as bowel perforation, urinary tract infection, and aspiration pneumonia ^5–7^. As the worldwide population ages, there is increasing need to establish a mechanistic understanding of the sepsis-aging paradigm, allowing for the development of innovative therapeutic strategies personalized to the older population.

In the most simplistic terms, sepsis is a severe host-pathogen interaction. However, sepsis definitions (and research investigations) are largely focused on the host response to the pathogen, as opposed to the pathogen itself ^8^. Within this framework, the pathogen is often seen as a static, homogeneous infectious insult that triggers the dysregulated host response. Accordingly, exaggerated sepsis severity outcomes in the aging population have been attributed to either an age-associated waning of immune function (i.e., immunosenescence), or an alteration in baseline inflammatory response (i.e., inflammaging) ^9,10^. However, therapeutics targeting the host immune response or inflammatory cascade have consistently failed to improve clinical outcomes of septic patients, in any age group ^11,12^. Conversely, therapeutic strategies targeting the pathogen with antimicrobial agents have consistently demonstrated significant decreases in sepsis-associated morbidity and mortality ^13,14^.

Given the importance of the pathogen to sepsis outcomes, we sought to determine if longevity-associated changes in gut microbial virulence contribute to aging-associated sepsis severity. We hypothesized that throughout the lifetime of the host, the gut microbiota accumulate virulence factors that promote host immune evasion. Escape of these age-conditioned pathogens from the intestinal lumen therefore leads to exaggerated sepsis severity. This novel concept — that the gut microbiota also “ages” throughout the lifespan of the host and selects for hypervirulent pathobionts — has the potential to inform targeted therapeutic approaches to mitigate the burden of sepsis in older adults ^3,9,15^.

## RESULTS

### Gut-microbiota induced sepsis severity is determined by the age of animal from which the infectious insult was derived

To determine the relative contribution of longevity-associated changes in the gut microbiome to sepsis severity, we utilized two complementary experimental sepsis models. Cecal ligation and puncture (CLP), the gold standard model of experimental sepsis, induces physiologic sepsis via leakage of the animal’s own gut bacteria into the peritoneal space ^16,17^. Although not the primary goal of our study, we employed CLP to demonstrate divergent sepsis severity outcomes in young mice and aged mice at the 24 hour time point (**Fig. 1A**) and establish experimental relevance to sepsis outcomes in older humans. Similar to previously published reports ^18,19^, aged mice exhibit increased mortality (17% [2/12] vs 0% in sham control and young mouse CLP), worsened acute kidney injury (**Fig. 1B**), and elevated plasma levels of the inflammatory cytokine Interleukin-6 (**Fig. 1C**) after CLP. To assess the attributable role of the infectious insult to this phenotype, we leveraged a second model of gut microbiota-induced experimental sepsis — intra-peritoneal fecal slurry injection. In this model, cage stool is collected and suspended in sterile saline prior to centrifugation and injection ^20^. Importantly, the fecal slurry model allows us to directly compare the virulence of fecal microbiota collected from one mouse population (i.e., fecal slurry from cages containing young animals, “FS(Y)”) versus fecal microbiota collected from another mouse population (i.e., fecal slurry from cages containing aged animals, “FS(A)”).

**Fig. 1.**
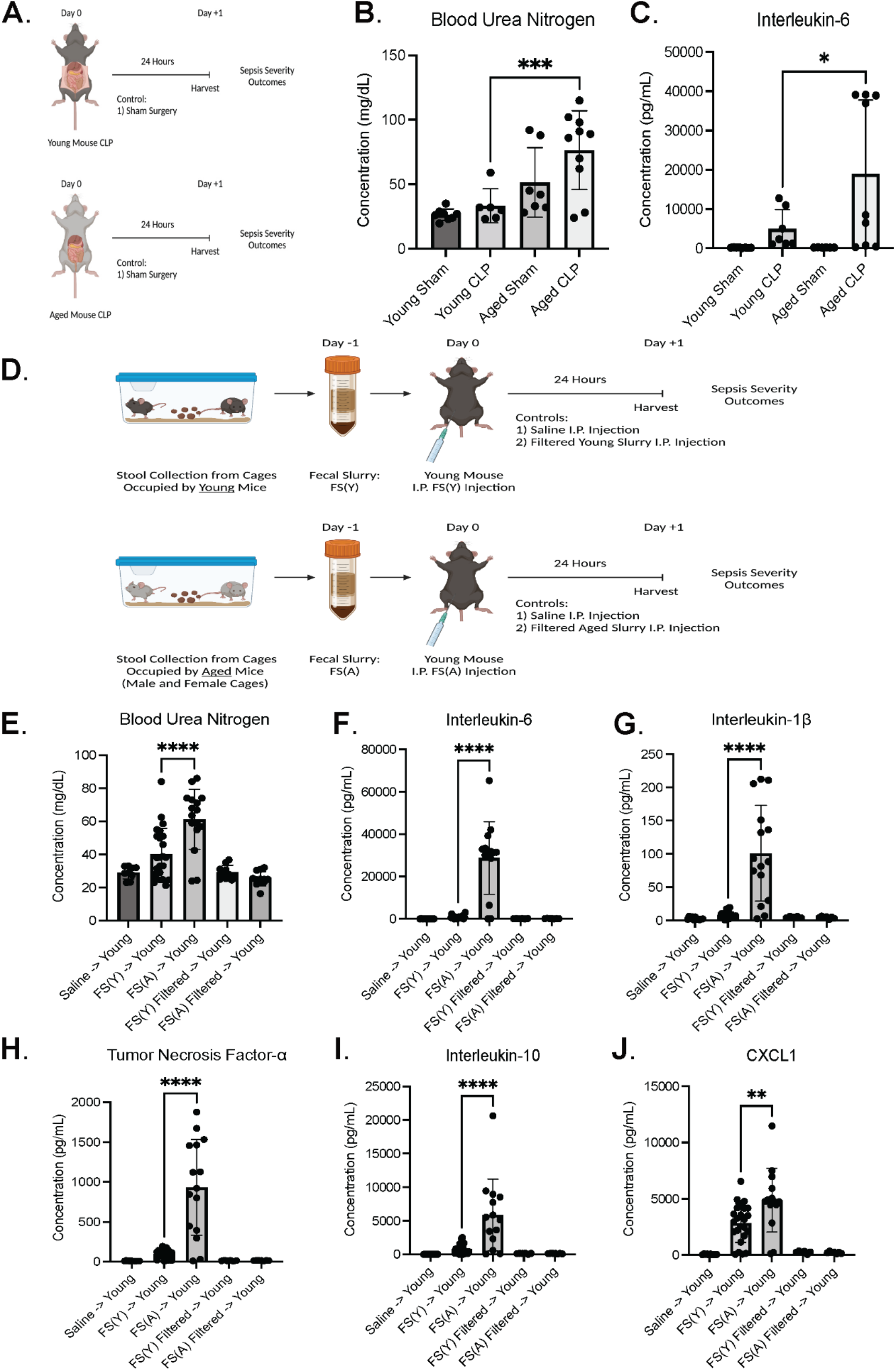
Age of animal from which infectious insult is derived determines sepsis outcomes. **(A)** Experimental design and measurements of sepsis severity via blood urea nitrogen **(B)** and circulating interleukin-6 **(C)** after cecal ligation and puncture compared between young and aged mice and with contemporaneous sham surgery control. *N* = 7-12 animals per group. Experimental dropout (i.e. mortality) 0% in all groups besides Aged CLP with 17% [2/12 animals] 24 hour mortality. **(D)** The complementary fecal slurry experimental sepsis model was utilized to assess the relative contribution of the aged gut microbiome to sepsis severity phenotype in young mice. Sepsis severity markers include acute kidney injury **(E)**, circulating plasma cytokines **(F-I)**, and chemokines **(J)** 24 hours after intraperitoneal fecal slurry injection. Intraperitoneal injection of saline and filtered slurry (0.22 micron filter) served as control conditions. *N* = 10-25 animals per group. Experimental dropout was 0% in all groups besides aged slurry (FS(A)) injection into young mice (32% mortality, [8/25 animals]). * = *p* < 0.05, ** = *p* < 0.01, *** = *p* < 0.001, **** = *p* <0.0001. Schematics (**Fig 1A, 1D**) created with BioRender.com.

This novel experimental design allowed for characterization of the gut microbiome as a contributor to aging-associated augmentation of sepsis severity. Our primary model kept the age of the recipient animal constant (young) while altering the donor age of the infectious insult (FS(Y) vs FS(A)) (**Fig. 1D**). Intriguingly, we found that the exaggerated sepsis severity observed in aged mice undergoing CLP could be replicated in young mice by inducing experimental sepsis with aged microbiota (FS(A)) as measured by 24 hour mortality (32% [8/25] vs 0% in FS(Y) and control groups), septic kidney injury (**Fig. 1E**), and plasma cytokine/chemokine response (**Fig. 1F–1J**). FS(A) induced the same augmented sepsis severity when administered to aged mice, with 100% 24 hour mortality [4/4] after injection with FS(A) and 0% 24 hour mortality [0/4] post FS(Y) injection. To determine if live bacteria were responsible for fecal slurry-induced sepsis, we filtered slurry through a 0.22 micron filter prior to injection. Mice receiving filtered slurry (either FS(Y) or FS(A)) experienced no mortality at 24 hours, no kidney dysfunction, and no elevation of plasma cytokine levels compared to saline control animals (**Fig. 1E–1J**).

To confirm that our findings were not simply an artifact of a unique batch of mice, we repeated these fecal slurry experiments using aged animals from multiple different sources including the National Institute on Aging and animals aged at two different institutions in Colorado, USA (University of Colorado Anschutz Medical Campus and Denver Health Medical Center). Similar findings were observed when fecal slurry was created from cages containing aged female mice (**Fig. 1D–1J**), suggesting that the increased virulence of FS(A) was not influenced by biological sex. Using both culture-dependent and culture-independent methods we demonstrated no differences between FS(A) and FS(Y) in the quantity of stool microbiota (**Fig. S1**).

### Longevity is not associated with an overabundance of known pathogenic taxa in the murine gut microbiome

Our phenotypic data led us to determine if there were differences in the overall composition of young versus aged murine gut bacterial communities. Utilizing 16S rRNA gene sequencing and analysis, we profiled microbiota present in the fecal slurry created from the two age groups. Beta diversity measures as represented by relative abundance (**Fig. 2A**) and principal component analysis (**Fig. 2B**) demonstrate an age-associated separation between groups (*p* = 0.027), consistent with previously published reports ^21–24^. There was a statistically significant increase in richness of aged stool slurry (*p* = 0.030), but no difference in evenness and overall alpha diversity (Shannon H index) (**Fig. 2C**). Only 5 genus-level taxa were more abundant in young stool slurry versus aged (with a fold-change > 2.0 and *p* < 0.05), conversely, there were no overabundant taxa identified in aged stool slurry (**Fig. 2D, 2E**). Taken together, these data suggest that while there are expected differences in microbiota associated with the aging process, the increased virulence of FS(A) was not simply due to an overabundance of known pathogens.

**Fig. 2.**
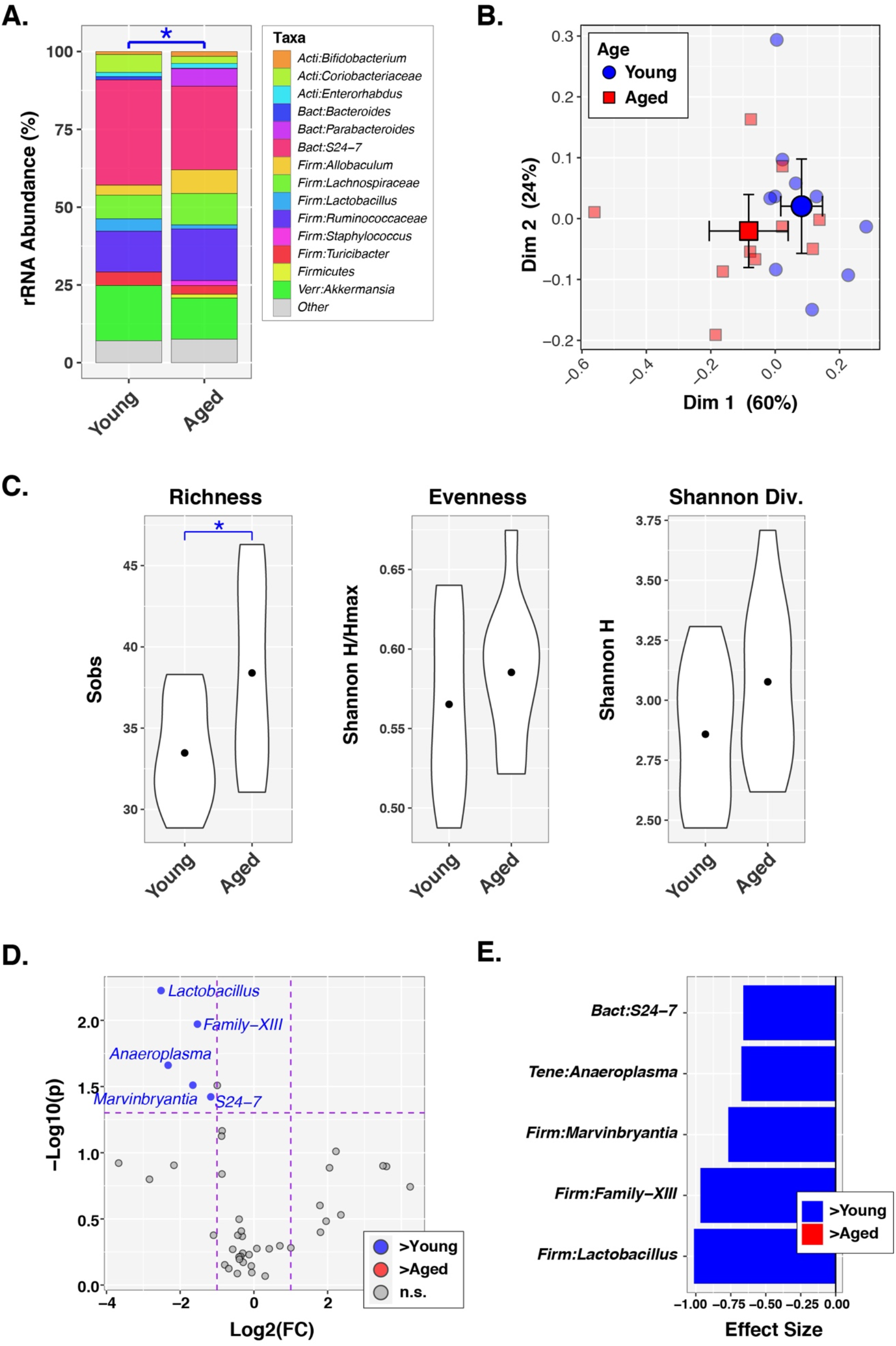
Longevity is not associated with an overabundance of known pathogenic taxa. Fecal slurry utilized for in vivo experiments subsequently underwent 16S rRNA gene sequencing and analysis for description of bacterial communities. *N* = 10 contemporaneous pairs of FS(Y) and FS(A) which is created from cages containing 5 animals per cage. **(A)** Relative abundance chart summarizing microbiota distributions between age groups. * = *p* <0.05 measured by a PERMANOVA test. **(B)** Principal coordinates analysis plot showing individual slurry samples (small circles, squares) and group means with 95% confidence intervals (large circles, squares). **(C)** Alpha diversity indices represented by violin plots indicating mean values (small circles) and distribution of values. * = *p* < 0.05 as measure by ANOVA. **(D)** Volcano plot demonstrating differentially abundant taxa in young (blue) and aged (red) mice, as determined by cutoffs of *p* value <0.05 and log-fold change > 2.0. **(E)** Effect sizes for individual significant taxa from volcano plot.

### Aging correlates with an overabundance of bacterial virulence factors present in the murine gut microbiome

The relatively subtle differences in microbial community composition observed in FS(Y) and FS(A) did not readily explain the divergent sepsis severity outcomes noted in our experimental sepsis models (**Fig. 1**). To further investigate these communities, we pursued whole genome sequencing and metagenomic analysis of the same fecal slurry samples used for in vivo experiments and 16S rRNA gene sequencing, in order to determine differences in the overall virulence capability of the bacterial communities between age groups. We utilized the Virulence Factor Database (VFDB) ^25^ to quantify the number and category of bacterial DNA sequences aligned against known proteins associated with virulence present in each group. **Fig. 3A** presents these data in volcano plot format, with heatmap and specific genes (and associated COG ^26^ [Cluster of Orthologous Genes] annotation in parentheses) coded for general virulence category (**Fig. 3B**). These findings demonstrate that FS(A) was enriched in bacterial genomes carrying more predicted virulence factors than FS(Y). This longevity-associated increase in microbiome virulence genes was driven by overrepresentation of exopolysaccharide, chemotaxis, flagella, and siderophore genes (**Fig. 3B**).

**Fig. 3.**
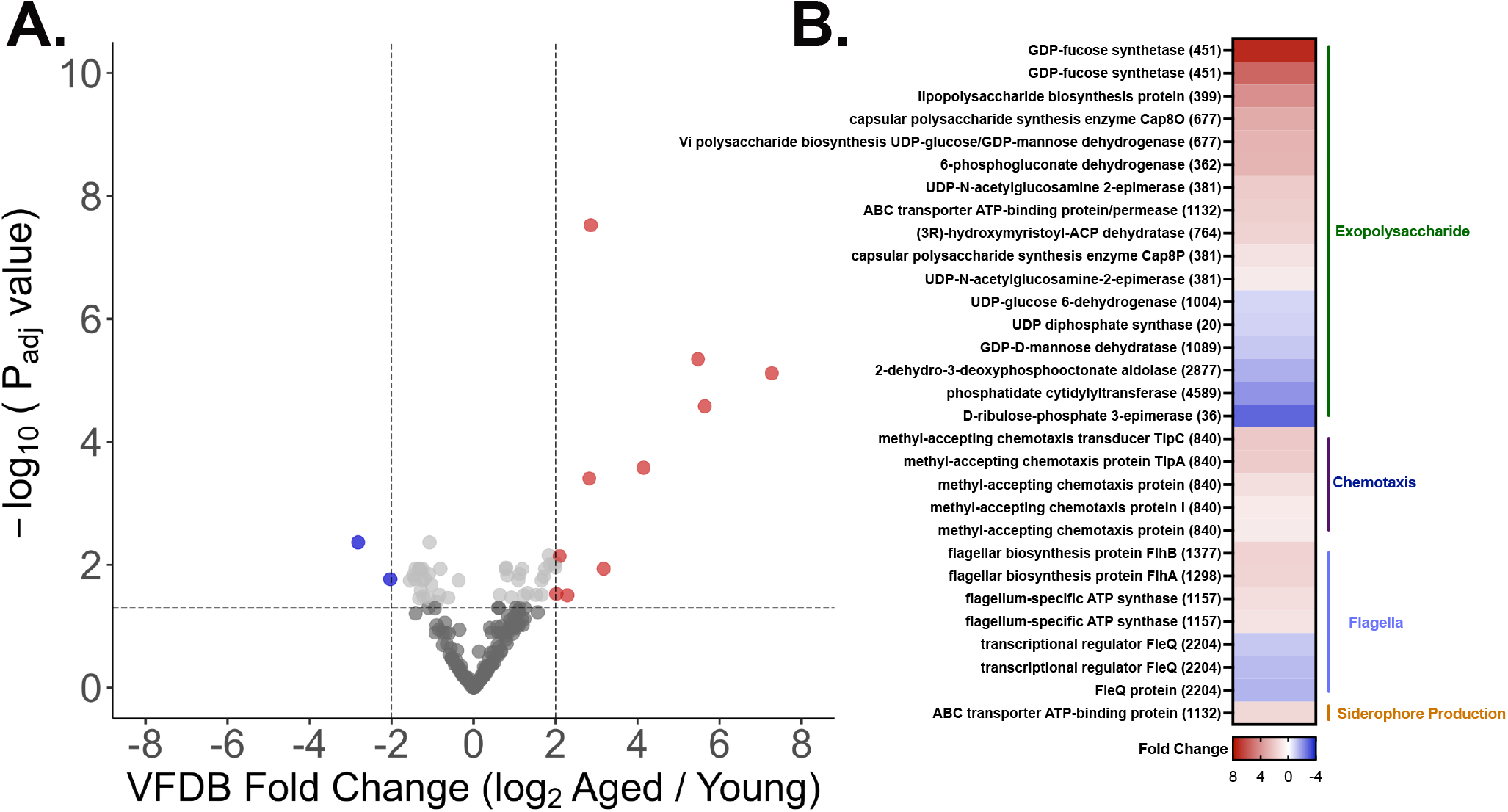
Aging is associated with an overabundance of virulence factors in the gut murine microbiome. The same fecal slurry samples with high quality bacterial DNA (FS(Y) *n* = 8 and FS(A) *n* = 6) from in vivo experiments and 16S rRNA analysis underwent whole genome shotgun sequencing and targeted metagenomic analysis focused on virulence factor abundance utilizing the Virulence Factor Database (VFDB) ^25^. **(A)** Volcano plot showing VFDB hits overabundant in FS(A) (red circles) versus FS(Y) (blue circles). Cutoff values of *p* <0.05 and log-fold change > 2. **(B)** Heatmap with all statistically significant (*p* <0.05) individual virulence factor genes and relative fold-change coded by color.

### Aging is associated with selection for pathogens resistant to blood killing in the murine gut microbiome

To determine if this longevity-associated enrichment in microbiome virulence had functional consequence, we focused further investigation into genes identified in FS(A) compared to FS(Y) which may explain in vivo sepsis severity alterations. We found that the aging stool microbiome is enriched with an overabundance of virulence factors associated with potential host immune evasion strategies including exopolysaccharide synthesis and maintenance (**Fig. 3B**). To determine the functional impact of these genetic differences on host immune evasion, we exposed FS(Y) and FS(A) to murine whole blood for 1 hour to allow for host mediated killing, followed by growth in media for 18-24 hours prior to plating and quantification of surviving colonies (**Fig. 4A**). Strikingly, no FS(Y)-derived bacteria survived blood killing, while multiple FS(A)-derived phenotypic colonies survived at high quantities after exposure to murine whole blood (either from young mice or aged mice, **Fig. 4B**, **Fig. S2**). Using MALDI-TOF (Matrix Assisted Laser Desorption/Ionization – Time of Flight) mass spectrometry, we identified that an *Escherichia coli* and an *Enterococcus faecalis* isolate (derived from FS(A)) survived whole-blood killing. Our findings suggest that the exaggerated sepsis severity noted in aged mice may be explained by a longevity-associated acquisition of host immune resistance genes in the gut microbiome.

**Fig. 4.**
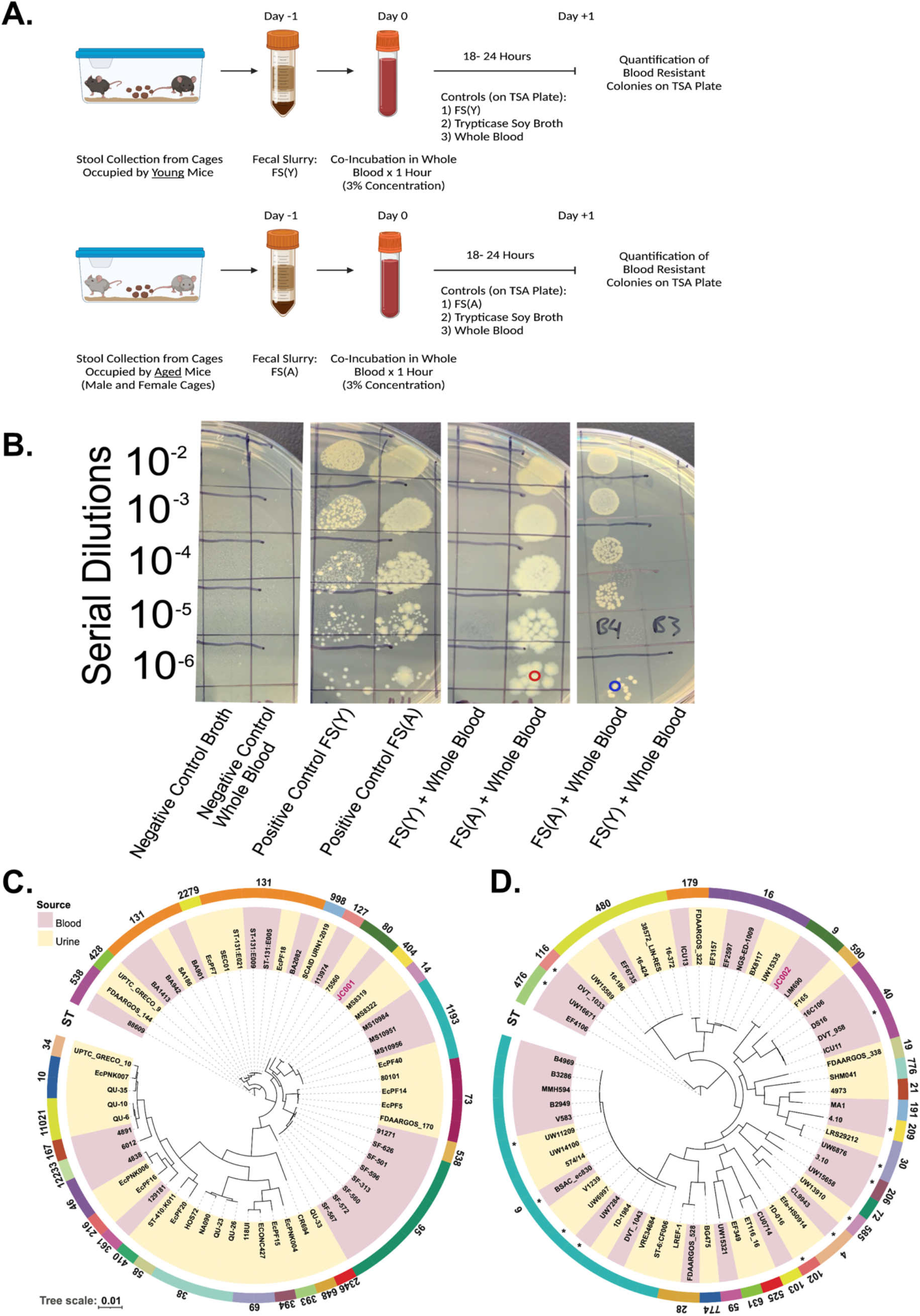
Aging selects for blood resistant pathogens in the murine gut microbiome. **(A**) Study design demonstrating screening process for whole blood killing resistance in fecal slurry. Biologic replicates were performed using young mouse blood, and aged mouse blood (**Fig. S2**). Control conditions included trypticase soy broth (negative control), whole blood (negative control), and FS(Y) and FS(A) without co-incubation with blood (positive control). After 1 hour of co-incubation in blood samples were allowed to grow overnight followed by serial dilution and quantification of colonies. **(B)** Representative image showing all control conditions and two experimental replicates of FS(Y) + Whole Blood and FS(A) + Whole Blood. Blood resistant colonies were identified by MALDI-TOF; red-circled colony = *E. coli* (JC001), blue-circled colony = *E. faecalis* (JC002), followed by whole genome sequencing **(C, D).** Core genome phylogeny of *E. coli* and *E. faecalis* isolates from human blood and urine. Maximumlikelihood trees of publicly available *E. coli* (*n* = 59) and *E. faecalis* (*n* = 58) isolates as determined by Roary and RAxML along with JC001 and JC002 isolated in this study. Isolates are colored as per their source of origin indicated in the key (red = blood, yellow = urine). The phylogenetic tree was visualized and annotated using iTOL. An asterisk (*) denotes that the sequence type (ST) was inconclusive, and the depicted ST is the nearest ST determined. Schematic **(A)** created with BioRender.com.

### Similar bacterial genetic features are seen in clinical bacterial isolates and the aged human gut microbiome

To determine whether our murine findings are relevant to increased sepsis severity in aged humans, we performed in-depth genetic analysis of our identified blood resistant bacterial isolates by comparative genomics. *E. coli* and *E. faecalis* are both frequent human pathogens and known inhabitants of the human gut microbiota. We sequenced these bacterial survivors of whole-blood (**Fig. 4**) and annotated their complete genomes (both chromosomal and non-chromosomal elements, **Data. S1**) in reference to the VFDB ^25^. Figures **4C** and **4D** present phylogenetic trees of these organisms and their relatedness to human-derived clinical isolates from blood or urine samples (additional details regarding the clinical isolates are presented in **Data. S2**). The isolated *E. coli* belongs to phylo-group B2, which is associated with human disease and is closely related to multiple additional clinical isolates ^27^. The *E. faecalis* isolate was identified as sequence type 9, which has documented clinical relevance and relatedness to known clinically virulent strains ^28^.

To further investigate the clinical relevance of our findings, we analyzed data from a previously published human microbiome metagenomic study to assess if there is a global shift towards aging-associated overabundance of virulence genes ^29^. Rampelli et al. generated fecal metagenomic data from 4 groups of patients with mean ages of 32.2, 72.5, 100.4, and 106.3 years respectively. We combined the two oldest groups into one “Aged Human” group (*n* = 38 patients total) and performed the same analysis pathway from our murine studies on these data with the VFDB ^25^. The volcano plot and heatmap of individual genes (with COG annotation) demonstrate an overabundance of gut microbiome virulence genes associated with human aging (**Fig. 5**). Analogous to our studies of gut microbiome virulence associated with murine aging (**Fig. 3**), the over-represented genes are classified as belonging to the same four mechanistic categories. The aged human microbiome was enriched in siderophore production genes, with a notable expansion of the number and fold change of yersiniabactin biosynthetic protein (COG0500) genes (**Fig. S3**) compared to aged murine counterparts.

**Fig. 5.**
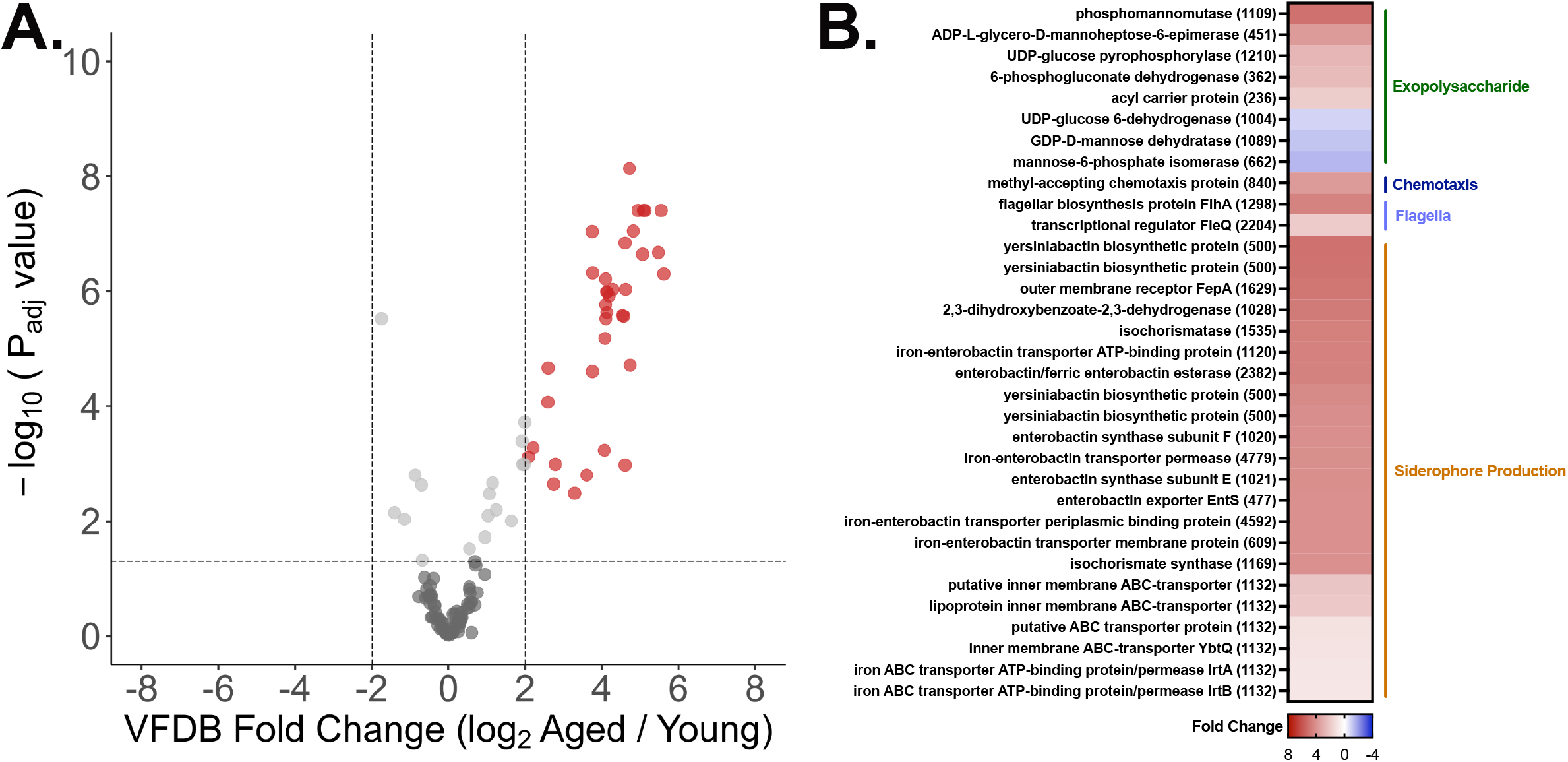
The aging human gut microbiome demonstrates an analogous overabundance of genomic virulence factors. Data from a previously published metagenomic dataset ^29^ underwent the same analysis strategy as our murine data (Fig 3). *n* = 62 total patients, *n* = 38 in “Aged Human” group. **(A)** Volcano plot showing VFDB ^25^ hits overabundant in aged human gut microbiome (red circles) versus young human gut microbiome (blue circles). Cutoff values of *p* <0.05 and log-fold change > 2. **(B)** Heatmap with all statistically significant (*p* <0.05) individual virulence factor genes and relative fold-change coded by color.

### Exopolysaccharide genes overabundant in the aging microbiome promote blood survival

One benefit of COG annotation is a de-duplication of similar genes encoding similar proteins in various prokaryotic organisms. The COG database (NCBI) identifies 4,877 COGs associated with over 3 million protein IDs. Analysis of virulence genes present in the aged murine gut microbiome, aged human gut microbiome, blood resistant *E. coli*, and blood resistant *E. faecalis* demonstrated thematic convergence onto exopolysaccharide pathways and three specific COGs (**Fig. 6A, 6B**). All three of these COGs are associated with exopolysaccharide synthesis and could conceivably explain resistance to host killing via various mechanisms (**Fig. 6C**). As a proof-of-concept experiment, we genetically manipulated one of the identified, overlapping COGs (COG0451) in a clinical strain of *E. faecalis* (V583, a human bloodstream isolate). The enterococcal polysaccharide antigen (Epa) operon has previously been described and noted to influence evasion of phagocytic killing ^30^, resistance to cationic antimicrobial peptides ^31^, and susceptibility to bacteriophage infection ^32^. We first utilized an engineered knockout of an Epa variable gene (*epaAC*, belonging to COG0451) in the V583 background. We conducted whole blood killing experiments with wild type V583, the V583 *epaAC* knockout strain (Δ*epaAC*), and a plasmid complemented Δ*epaAC* strain to test the functional importance of this gene for blood survival. **Fig. 6D** shows that the V583 Δ*epaAC* strain was more susceptible to blood killing than wild type V583, and that resistance to blood killing could be restored by plasmid complementation of the *epaAC* gene. In addition to this engineered knockout strain, in a screen of bacteriophage resistant *E. faecalis* (V583), we identified a naturally occurring strain (4RSR) with a missense point mutation in the *epaAC* gene, resulting in a V84L substitution. Similarly, this strain demonstrates markedly reduced blood survival presumably due to a nonfunctional Epa product (**Fig. 6E**).

**Fig. 6.**
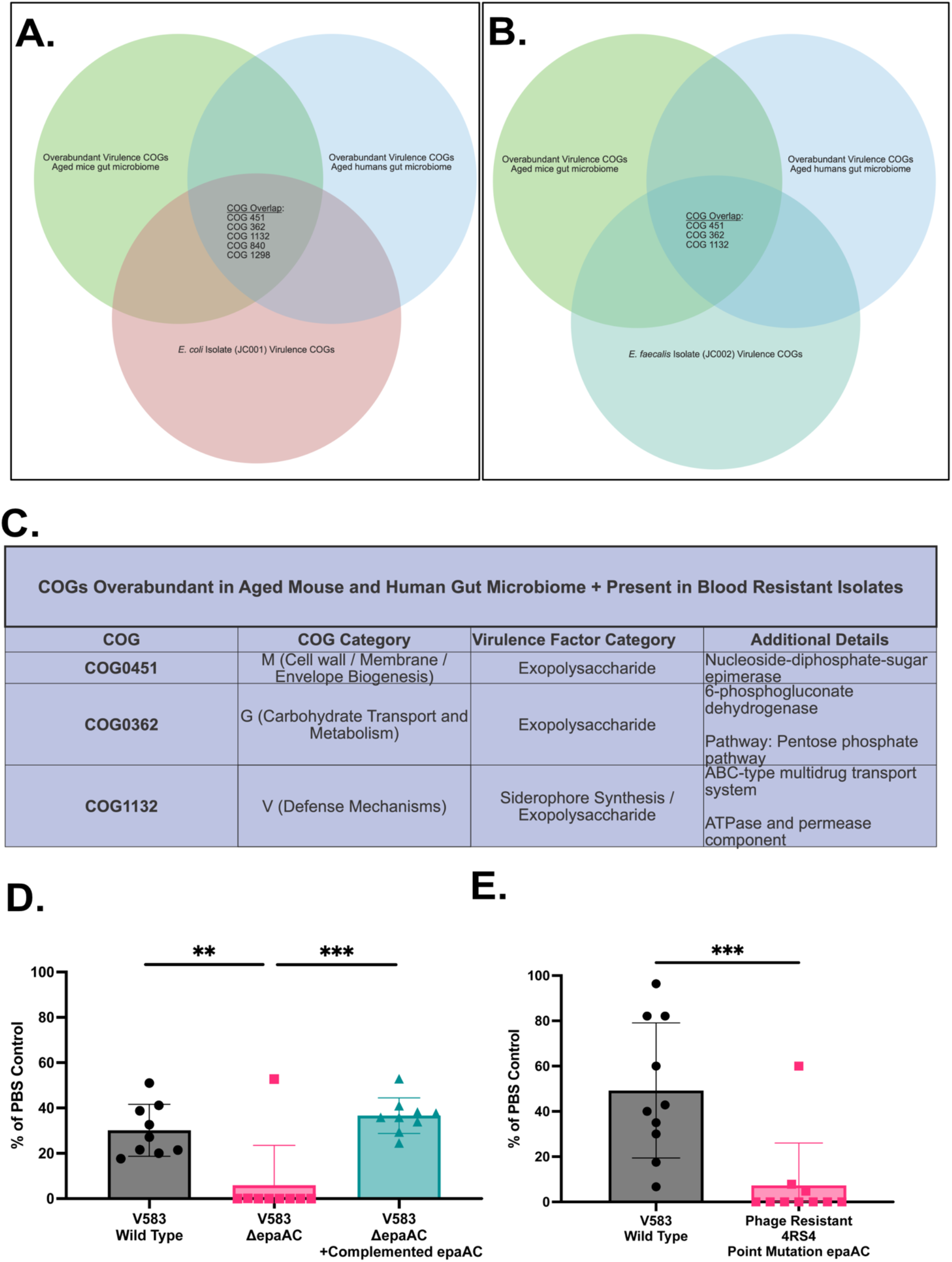
Identified exopolysaccharide virulence genes promote blood survival. **(A)** Venn diagram demonstrating Cluster of Orthologous Gene (COG) overlap between the aged murine gut microbiome, aged human gut microbiome, and isolated (blood-resistant) *E. coli*. **(B)** Analogous Venn diagram with isolated, blood-resistant *E.faecalis*. **(C)** Table highlighting in detail the three COGs overabundant in both murine and human gut microbiome, as well as both blood-resistant isolates. **(D,E)** Blood killing assay of various strains of *E.faecalis* with genetic manipulation of the enterococcal polysaccharide antigen (*epaAC*) gene (COG0451). Data presented as percent of PBS control (no blood killing). **(D)** Comparison of wild type (V583 isolate) *E.faecalis* blood survival versus an engineered *epaAC* knockout and *epaAC* knockout + plasmid complementation. (**E)** Comparison of V583 versus a phage-resistant strain (4RSR) with a missense point mutation in *epaAC* rendering the product non-functional. *N* = 9 per group (3 biologic replicates and 3 technical replicates per group). Paired *t* test statistical analysis with * = *p* < 0.05, ** = *p* < 0.01, and *** = *p* < 0.001.

## DISCUSSION

Our findings highlight a previously unrecognized contributor to the pathophysiology of heightened aging-associated sepsis severity. To date, investigations of the intersection of aging and critical illness have focused on longevity-associated host processes such as waning immune function and alterations in inflammation. However, it is intuitive that the intestinal microbiota simultaneously undergoes genomic and phenotypic changes throughout the lifespan of the host organism. This aging of an enteric bacterial community likely selects for pathobionts with virulence factors that offer a fitness benefit — such as the evasion of host immunity — as demonstrated ex vivo by resistance to whole blood killing (**Fig. 4**). Our work highlights that pathogen virulence factor genomics, and not simply type of pathogen, is therefore a major mediator of mechanistic sepsis heterogeneity.

Importantly, this concept has the potential to inform the pursuit of novel biomarkers and therapeutic targets. The inherent complexity of these biologic systems makes the discovery of a single unifying mechanism highly improbable — indeed, a single virulence factor is unlikely to fully explain the entirety of aging-associated sepsis severity. Our virulence-focused metagenomic analyses identified numerous overabundant, age-associated virulence factors in the gut microbiome from both mice and humans. However, it is important to note that the individual genes were not identical between host species. We pursued mechanistic investigation into COG0451 due to the presence of this COG in blood resistant isolates, and both aged human and murine microbiomes (**Fig. 6**), but there were additional examples of remarkable genomic overlap. Multiple siderophore biosynthetic genes were found to be overabundant in aged humans and homologous biosynthetic clusters were found in the *E. coli* isolate (JC001) recovered from aged murine stool (**Fig. S3**). One of these siderophore production operons produces yersiniabactin (Ybt), a siderophore implicated in the pathogenesis of multiple gram-negative species ^33–35^. Ybt functionality increases *E. coli* virulence and has been shown to both increase availability of nutritional iron and protect against iron toxicity through sequestration mechanisms ^35,36^. The operon encoding for Ybt is contained in the *Yersinia* high-pathogenicity island, a mobilizable genomic component ^37^. The presence or absence of the various identified microbial genomic characteristics may provide the basis for accurate risk stratification and alternative antibacterial therapeutics designed to restore bacterial susceptibility to host immune factors.

The forces leading to this the microbial shift towards enhanced virulence capability with aging remain unexplored. We speculate that a slow evolutionary selective pressure associated with repeated host-pathogen interactions at the gut mucosal barrier inform these changes in a stochastic fashion. However, the spread of mobilizable genomic features such as the previously discussed *Yersinia* pathogenicity island in the aged microbiota provides an alternative hypothesis for elevated longevity-associated virulence potential. It is also important to consider that this phenomenon may not be unique to aging; there may be analogous changes in other disease states, behaviors, or iatrogenic factors that predispose to exaggerated sepsis severity via a gut microbiota with an increased armament of virulence factors. This construct has the potential to better explain the heterogeneity of human sepsis outcomes based on underlying comorbidities, therapies, diet, and other factors which requires further investigation.

There are more proximal potential clinical applications for the presented concept as well. A major clinical challenge is the prevention of secondary nosocomial infection in the intensive care unit either post primary infection, or post non-infectious diagnosis. To address this concern, prior work has explored selective decontamination of the digestive track with broad spectrum, enteral, antibiotic therapy ^38^. While clinical benefit of this strategy has been demonstrated, it has not been widely adopted in the United States due to logistical hurdles and concern for the selection of multidrug resistant organisms. This project lays the framework for sophisticated risk assessment of the virulence capacity of the gut microbiome and resulting targeted decontamination therapy to prevent nosocomial infection syndromes on an individual patient basis. Although speculative, our data suggest that directed bacteriophage therapy (as an alternative to broad spectrum antibiotic therapy) may select for phage-resistant — but as a tradeoff — host-susceptible pathogens in the gut microbiome (**Fig. 6**).

Our work has additional, unanticipated implications for pre-clinical sepsis research and experimental sepsis models utilizing gut-derived bacteria (i.e., cecal ligation and puncture, CLP, the gold standard model of experimental sepsis). In CLP, mice are exposed to their own gut microbiota. Our findings suggest that differences in CLP outcomes between different mouse genotypes may not merely be a function of host phenotype, but also may reflect differential accumulation of virulence factors in enteric bacteria. It is thus plausible that microbial changes are driving outcomes in various murine models including pre-treated animals and transgenic strains. We encourage other researchers to take this variable into account in their mechanistic sepsis investigations, potentially by complementing CLP with fecal slurry modeling, using stool collected from a shared mouse donor.

Finally, our study creates a translational rationale supporting future prospective clinical studies. In our murine models, we have demonstrated an association of increased microbiome virulence with both age and with sepsis severity outcomes. While we note a similar increase in microbiome virulence and striking overlap of abundant virulence factors in the gut microbiome from older humans, additional prospective studies will be necessary to determine if this increase in bacterial virulence translates to worsened sepsis outcomes in aged humans. Another limitation of this work is a focus on bacterial aspects of the gut microbiome, although we acknowledge that fungal, archaeal, and viral components are also present in the mammalian gut microbiota. Due to a paucity of available techniques and lack of obvious aging-associated clinical relevance, we have not pursued investigations into these additional kingdoms of life in the current study.

In summary, our work identifies that aging-associated sepsis severity not only reflects host factors, but also may be driven by the effects of longevity on pathogen virulence. These novel findings have major implications for our understanding of the mechanistic heterogeneity of critical illness, potentially guiding the personalization of sepsis care for the aged adult.

## ONLINE METHODS

### Study Design

The overall objective of this research was to assess longevity-associated alterations in the gut microbiota and their relative contribution to sepsis severity. All animal experiments were conducted under approved University of Colorado IACUC (Institutional Animal Care and Use Committee, Protocol #00307) and IBC (Institutional Biosafety Committee) protocols and in accordance with the ARRIVE (Animal Research: Reporting of In Vivo Experiments) guidelines ^39^. Individual animals were blindly randomized to experimental or control groups. Rigorous contemporaneous controls were used, as described in individual experiments. For our primary experimental sepsis model (fecal slurry injection) we performed a power calculation, based on prior work, and estimated 10-12 animals per experimental group to detect a 50% difference in sepsis severity (continuous variables; acute kidney injury and plasma cytokine levels) with a power of 90% and α of 0.05. However, due to our higher than anticipated mortality (approximately 1/3 in FS(A) injected animals), we doubled the number of animals in this experimental condition to account for drop out and to include biologic sex (FS(A) created from aged female cages) as a variable. Statistical outliers were defined prior to experiments (Graphpad Prism ROUT method with recommended Q value of 1%) and were removed prior to analysis. Animal drop-out (e.g. mortality) was recorded and explicitly reported. The intricacies of study design for specific experiments are presented in individual figures and figure legends.

### Animals

Young male (8-10 weeks of age) C57BL/6 mice were obtained from Jackson Laboratory (Bar Harbor, ME) and allowed to acclimate in our local vivarium for 1 week prior to experimentation. As discussed in the results section, we were concerned regarding the potential confounder of a “batch effect” of older animals so obtained aged animals (and stool) from various sources. Aged C57BL/6 male and female animals were acquired from the National Institute on Aging (Bethesda, MD) and allowed to acclimate for 1 week in Aurora, CO prior to experimentation. Additionally, C57BL/6 mice born and aged in in Denver, CO (at Denver Health Hospital, an academic partner of the University of Colorado) were utilized. Aged animals were used for experiments between the ages of 20-24 months. All animals were housed in the same vivarium room with identical diet, day/night cycle, and cleaning schedules.

### Experimental Sepsis Models

As previously described, we have utilized two complementary experimental sepsis models to isolate the role of the gut microbiome as a driver of outcomes ^20^.

#### Cecal Ligation and Puncture

Under inhaled isoflurane anesthesia, laparotomy was performed followed by visualization and isolation of the cecum. The cecum was ligated with a 2-0 silk suture and punctured once through-and-through with a 23 gauge needle. The punctured cecum was returned to the abdominal cavity and the incision was closed with 4-0 silk sutures. 500 μL of subcutaneous normal saline resuscitation was given. Analgesic subcutaneous buprenorphine and local bupivacaine were utilized. Sham surgery was identical to CLP, without ligation or puncture of the cecum ^20^.

#### Fecal Slurry Injection

Dry fecal pellets were collected one day before injection and suspended in sterile saline at a constant weight to volume ratio (180 mg stool per 1 mL of saline) and vortexed for 5 minutes. After storage overnight at 4°C, the sample was centrifuged (100 rpm x 5 minutes), and the liquid portion removed. 400 μL of fecal slurry was injected via the intraperitoneal route to induce sepsis. 400 μL of sterile saline injection was used as a contemporaneous control. Fecal slurry was also filtered through a 0.22 micron filter to remove live bacteria and serve as an additional control ^20^.

### Sepsis Severity Outcomes

#### Mortality

Animals were observed after induction of sepsis by either model. Animals identified as being moribund were euthanized and recorded as mortality at 24 hours.

#### Acute Kidney Injury

Plasma Blood Urea Nitrogen (BUN), an indirect measurement of glomerular filtration rate, was quantified 24 hours after sepsis induction using the Quantichrom Urea Assay Kit (BioAssay Systems, Hayward, CA). Working reagent from the kit was added to plasma samples and optical density was read to 520 nm after 20 minutes of reaction time.

#### Plasma Cytokine/Chemokine Quantification

Plasma Interleukin-1β, Interleukin-6, Tumor Necrosis Factor-α, Interleukin-10, and CXCL1 were quantified using Meso Scale Diagnostics (Rockville, MD) custom multiplex electrochemiluminescence detection arrays.

### 16S rRNA Microbiota Analysis

#### 16S rRNA Amplicon Library Construction

DNA was isolated from FS(Y) and FS(A) samples using the Qiagen PowerFecal DNA extraction kit. Bacterial profiles were determined by broadrange amplification and sequence analysis of 16S rRNA genes following our previously described methods ^40,41^. In brief, amplicons were generated using primers targeting the V3-V4 variable region of the 16S rRNA gene. PCR products were normalized using a SequalPrep™ kit (Invitrogen, Carlsbad, CA), pooled, lyophilized, purified and concentrated using a DNA Clean and Concentrator Kit (Zymo, Irvine, CA). Pooled amplicons were quantified using Qubit Fluorometer 2.0 (Invitrogen, Carlsbad, CA). The pool was diluted to 4nM and denatured with 0.2 N NaOH at room temperature. The denatured DNA was diluted to 15pM and spiked with 25% of the Illumina PhiX control DNA prior to loading the sequencer. Illumina paired-end sequencing was performed on the Miseq platform with versions v2.4 of the Miseq Control Software and of MiSeq Reporter, using a 600 cycle version 3 reagent kit.

#### Analysis of Illumina Paired-end Reads

Illumina Miseq paired-end reads were aligned to mouse reference genome mm10 with bowtie2 and matching sequences discarded ^42,43^. As previously described, the remaining non-mouse paired-end sequences were demultiplexed and then assembled using phrap ^44,45^. Pairs that did not assemble were discarded. Assembled sequences were trimmed over a moving window of 5 nucleotides until average quality met or exceeded 20. Trimmed sequences with more than 1 ambiguity or shorter than 350 nt were discarded. Potential chimeras identified with Uchime (usearch6.0.203_i86linux32) ^46^ using the Schloss ^47^ Silva reference sequences were removed from subsequent analyses. Assembled sequences were aligned and classified with SINA (1.3.0-r23838) ^48^ using the 418,497 bacterial sequences in Silva 115NR99 ^49^ as reference configured to yield the Silva taxonomy. Operational taxonomic units (OTUs) were produced by clustering sequences with identical taxonomic assignments. This process generated a median of 107,077 sequences/sample (IQR: 80505-136448). The software package Explicet (v2.10.5, www.explicet.org)^50^ was used to calculate alpha-diversity scores through 1000 replicate re-samplings. All sequence libraries had Goods coverage scores ≥ 99%. All 16S rRNA gene sequence data and associated metadata were deposited in NIH Genbank’s sequence read archive (SRA) under BioProject ID PRJNA838636.

### Metagenomic Analysis

#### BacterialDNA Reads

DNA was isolated from FS(Y) and FS(A) samples using the Qiagen PowerFecal DNA extraction kit and whole genome shotgun sequencing was performed by Novogene (Sacramento, CA). Human reads were obtained from the sequence resource archive (SRA) from accession number PRJNA553191 ^29^. Read decontamination and trimming were performed as previously described ^51^. In short, reads were trimmed using bbduk, part of the BBTools v38.90 bioinformatics tools using the following parameters: ktrim=l ktrim=r k=20 mink=4 minlen=20 qtrim=f ftl=20. Trimmed reads were mapped against human (hg38), mouse (mm39), and phiX174 genomes. Reads that did not map were used for the downstream analyses.

#### Mapping to the Virulence Factor Database

The core protein virulence factor database (VFDB) ^25^ used in this study was downloaded on June 28^th^, 2021. Decontaminated and trimmed reads were mapped against the database from both human and mouse samples using PALADIN v1.4.6 ^52^ with default settings. Total reads mapped to each virulence factor open reading frame were calculated using Samtools v1.13 ^53^. Raw counts were used as input for DESeq2 v 1.30.1 ^54^ to test for differential abundance, with zero-heavy rows eliminated from the count matrix before running the DESeq2 pipeline. All metagenomic gene sequence data and associated metadata were deposited in NIH Genbank’s sequence read archive (SRA) under BioProject ID PRJNA844935.

### Whole Blood Killing Assay

Murine whole blood was collected on day of experiment via inferior vena cava collection. For fecal slurry experiments (**Fig. 4**), slurry was created as previously described and co-incubated in whole blood at a concentration of 3% for one hour at 37°C. After incubation, 100 μL of trypticase soy broth was added to each well and the samples were allowed to grow overnight at 37°C. After 18-24 hours of growth serial dilution was performed for quantification of blood-resistant colonies. For individual *E. faecalis* bacterial isolates (**Fig. 6**), the strains were incubated in whole human blood (collected under Colorado Multiple Institutional Review Board protocol 17-1926). Blood killing assays were performed on *E. faecalis* V583, and isogenic derivatives of V583 including BDU62 (Δ*epaAC*, EF2165), BDU62 pPL05 (*epaAC* complement), and 4RS4 (*epaAC* Val 84 Leu mutant). *E. faecalis* V583 and its *epaAC* derivatives have been described previously ^32,55^. To determine the susceptibility of these strains to blood killing, each strain was grown overnight in Brain Heart Infusion (BHI) broth at 37°C with shaking and approximately 1×10^6^ CFU of each strain was added to either 90 μL of fresh human blood or phosphate buffered saline (PBS), mixed, and incubated statically at 37°C for 1 hour. 100 μL of BHI was added and the samples were incubated at 37°C overnight. Samples were serially diluted and plated on BHI agar and colonies were enumerated. Percent survival was calculated as the percent of cells surviving blood killing compared to control cells in PBS.

### Bacterial Identification by MALDI-TOF Mass Spectrometry

Fresh bacterial colony growth was deposited on a polished steel MSP 96 target (Bruker Daltonics, Leipzig, Germany) and covered with 1 μl of a 70% formic acid solution. Following air-drying, the bacterial spot was overlaid with 1 μl of a saturated α-cyano-4-hydroxycinnamic acid (HCCA) matrix solution (Bruker Daltonics). Mass spectra were acquired and analyzed using a Microflex LT mass spectrometer (Bruker Daltonics) in combination with research-use-only (RUO) versions of the MALDI Biotyper software (MBT Compass v4.1) and the reference database v.9.0.0.0 (8468 spectra covering 2969 species). Calibration was done by following the manufacturer’s instructions and using the manufacturer’s recommended bacterial test standard. Bacterial species were assigned for scores of ≥2.0.

### Comparative Genomics

#### Sequencing and assembly

The genomes of isolates JC001 and JC002 were sequenced and assembled by the Microbial Genome Sequencing Center (MiGS, Pittsburgh, PA). Genomic sequence data was deposited in NIH Genbank’s sequence read archive (SRA) under BioProject ID PRJNA844935 with individual accession numbers (JC001 = SAMN28860864, JC002 = SAMN28860875).

#### Genome annotation

Open reading frames were predicted and annotated using RAST-Tk ^56^. Contigs were assembled using megahit v1.2.7 with the “meta-large” preset ^57^ and open reading frames predicted and annotated using RAST-Tk with the determined annotations from the blood-killing isolates against metagenome contigs using BLASTn ^58^. Alignments between contigs were also performed using pairwise BLASTn. Contig visualization was performed using the gggenes package (v0.4.1) in R (v4.0.5).

#### Pan-genome analysis

Isolates JC001 and JC002 were analyzed with reference to publicly available genomic data of 59 human *E. coli* (blood, n=24; urine, n=35) and 58 human *E. faecalis* (blood, n=29; urine, n=29) isolates collected from the NCBI Genome database. Multi-Locus Sequence Typing (MLST) ^59^ was performed using the MLST Web server (available at www.cbs.dtu.dk/services/MLST) and the ‘*Escherichia coli* #1’ and ‘*Enterococcus faecalis*’ configurations. Genomes of all 119 isolates were annotated using Prokka ^60^. Core genome alignments were generated using the Roary ^61^ pipeline and were then used to construct maximum-likelihood (ML) trees using RAxML ^62^. iTOL ^63^ was used to visualize the phylogenetic trees and metadata.

#### E. coli phylo-typing

JC001 was determined to belong to *E. coli* phylo-group B2 by examining the genome for the presence of the *chuA*, *yjaA*, and the DNA fragment TSPE4.C2 and using the Clermont classification system ^64^.

### Statistical Analysis

For in vivo experiments, prior to analysis, statistical outliers were identified as prespecified using the ROUT method in Graphpad Prism with Q Value = 1%. Biologic and technical replicates (*n*) and drop-out (mortality) are noted in individual figure legends. Comparisons between multiple independent groups were conducted using an ordinary one way ANOVA (Analysis of Variance). Statistical significance of comparisons of interest (non-control conditions) was performed via Sidak’s multiple comparisons tests. Whole blood killing quantification statistical significance was assessed using paired *t* tests. A *P* value of < 0.05 was considered statistically significant.

For 16S rRNA gene sequence analysis, differences in overall composition (i.e., beta-diversity) were assessed through permutational ANOVA (PERMANOVA ^65,66^) with the Morisita-Horn dissimilarity index. PERMANOVA p-values were inferred through 10^6^ label permutations. Principal coordinates analysis was carried out using the vegan wcmdscale function. Alphadiversity indices (i.e., S_obs_, Shannon H, Shannon H/Hmax) were assessed by ANOVA. Individual taxa differing between treatment groups were identified using the ANOVA-like differential expression (ALDEx2) R package.^67,68^. Additional specifics of statistical strategy for genomic analyses are presented in individual subheadings above.

## Supplementary Materials

Fig. S1. There is no difference in total bacterial DNA between FS(Y) and FS(A).

Fig. S2. Donor age of whole blood does not impact pathogen blood resistance.

Fig. S3. Similar yersiniabactin operons are present in blood resistant *E. coli* isolate and overabundant in the aged human gut microbiome.

Table S1. Specific aging-associated differential virulence factors in the murine and human gut microbiome.

Data file S1. Virulence Factor Database annotation of complete blood resistant *E. coli* (JC001) and *E. faecalis* (JC002) genomes including COG number.

Data file S2. Comparison clinical *E. coli* and *E. faecalis* isolates utilized for comparative genomic and phylogenetic tree analyses.

## Funding

National Institutes of Health K08AG061144 (JFC)

National Institutes of Health R03AG056353 (JFC)

National Institutes of Health R01GM125095 (EPS)

National Institutes of Health R01AI141479 (BAD)

National Institutes of Health T32AI052066 (JMK)

National Institutes of Health R01AG018859 (EJK)

National Institutes of Health R01DK131267 (NJD)

Department of Veterans Affairs I01 BX002711 (ARH)

Department of Veterans Affairs Eastern Colorado GRECC (Geriatric Research, Education, and Clinical Center)

## Data and Materials Availability

All data are available in the main text or the supplementary materials. Genomic data and associated metadata were deposited in NIH Genbank’s sequence read archive (SRA) under the following BioProject ID’s:

- Murine Gut Microbiome 16S rRNA: PRJNA838636
- Murine Gut Microbiome Metagenomic Data: PRJNA844935
- Individual Bacterial Isolates: PRJNA844935

- JC001 (*E. coli*) = SAMN28860864
- JC002 (*E. faecalis*) = SAMN28860875

---

**Fig. S1.**
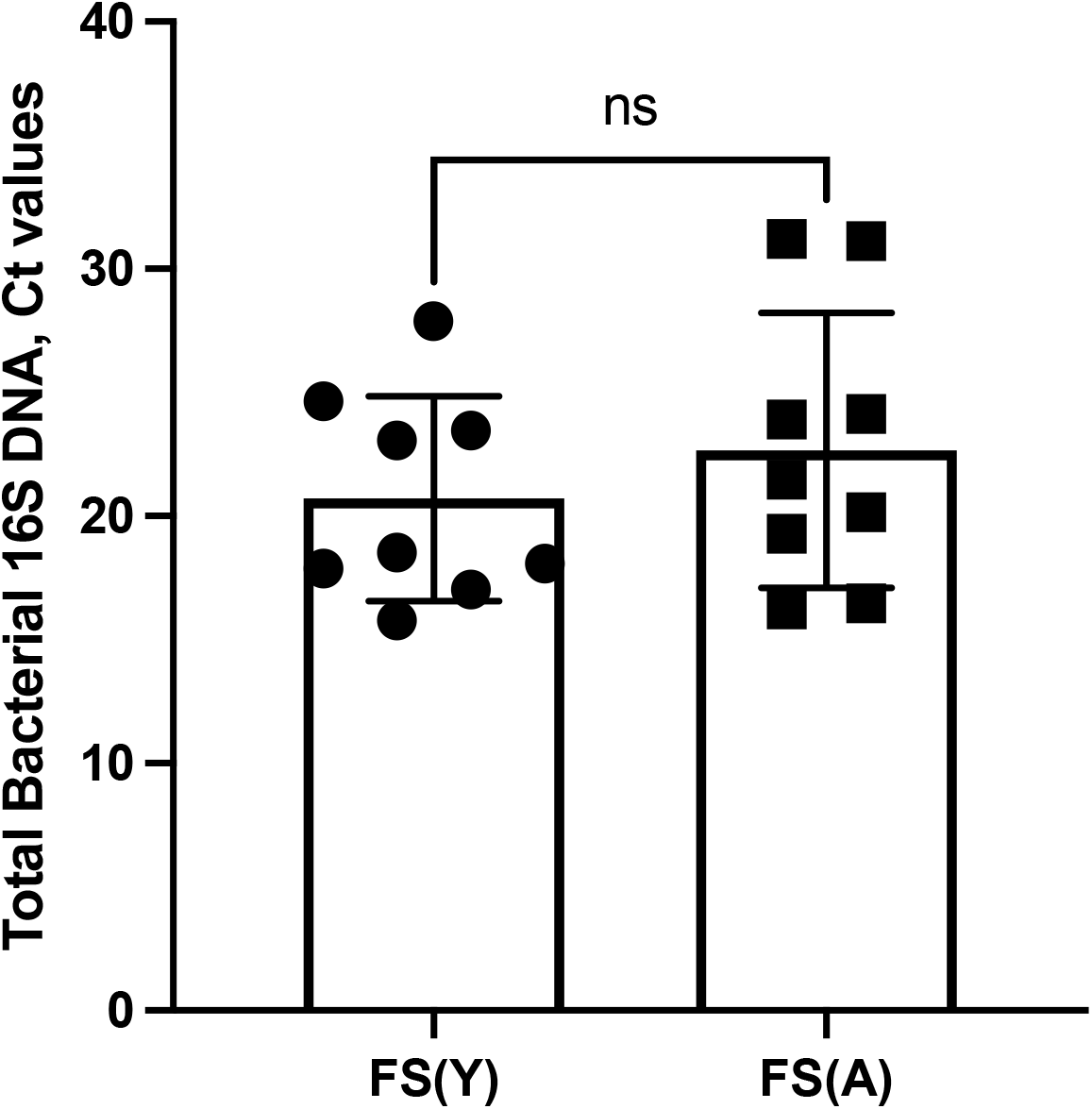
There is no difference in total bacterial DNA between FS(Y) and FS(A). Total bacterial 16S DNA was sequenced and these data are presented as cycle threshold (Ct) values. *N* = 9 contemporaneous pairs of FS(Y) and FS(A).

**Fig. S2.**
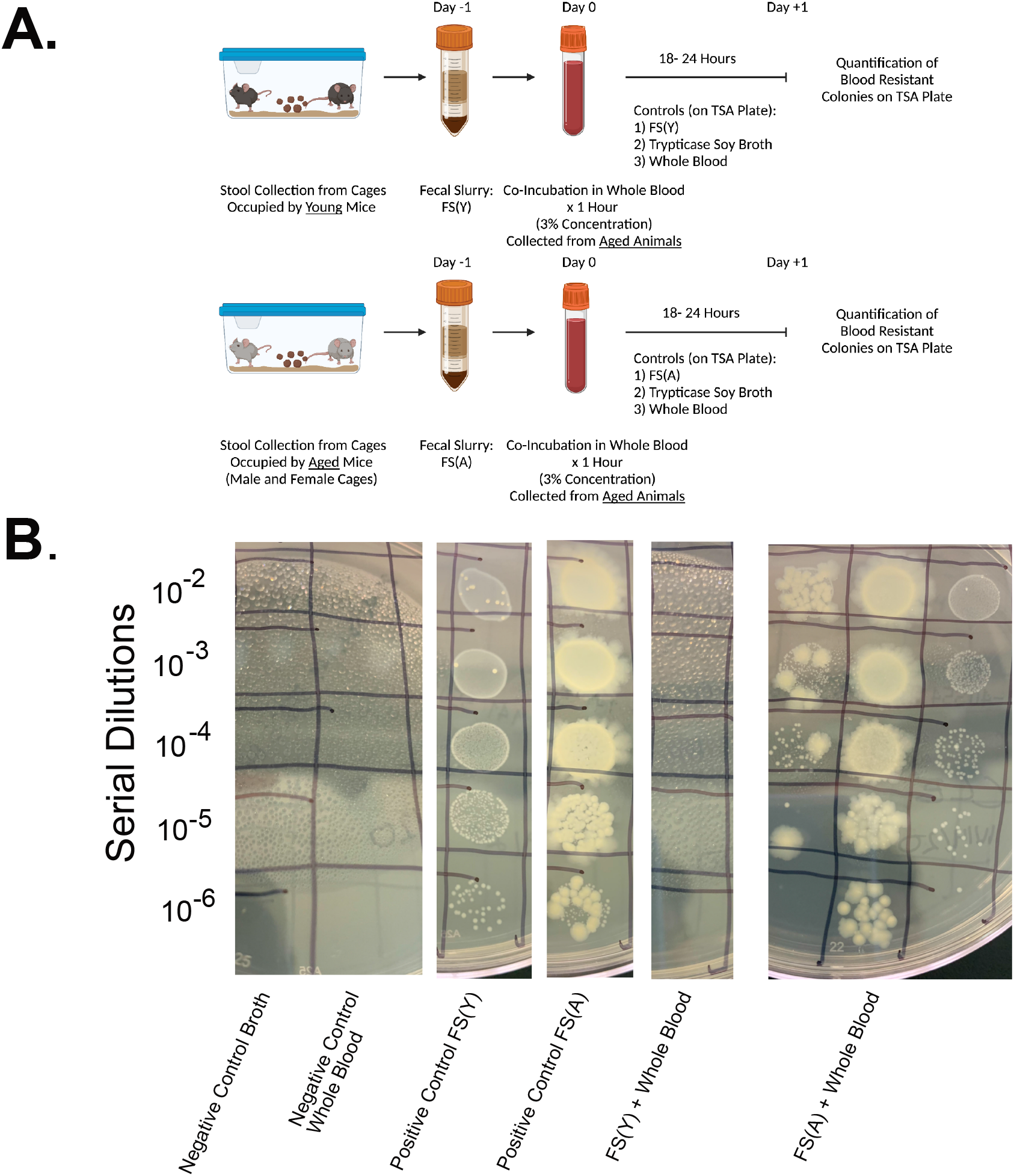
Donor age of whole blood does not impact pathogen blood resistance. **(A**) Similar to Fig. 4, study design demonstrating screening process for whole blood killing resistance in fecal slurry using aged mouse blood for the assay. Control conditions included trypticase soy broth (negative control), whole blood (negative control), and FS(Y) and FS(A) without co-incubation with blood (positive control). After 1 hour of co-incubation in blood samples were allowed to grow overnight followed by serial dilution and quantification of colonies. **(B)** Representative image showing all control and experimental conditions. Schematic **(A)** created with BioRender.com

**Fig. S3.**
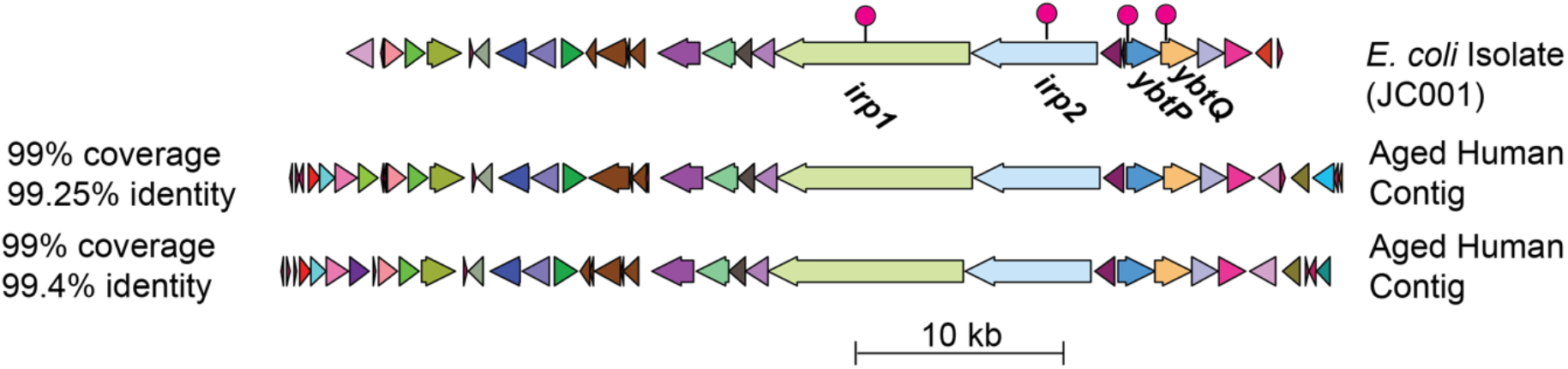
Similar yersiniabactin operons are present in blood resistant *E. coli* isolate and overabundant in the aged human gut microbiome. Comparison of yersiniabactin operons found in JC001 and aged human metagenomes. Genes labeled with lollipops and names were overabundant in aged human metagenomes. Color signifies similar gene function between operons. Coverage and identity between each aged human metagenome contig and the Ybt operon in JC001 was calculated with BLASTn.

**Table S1.**
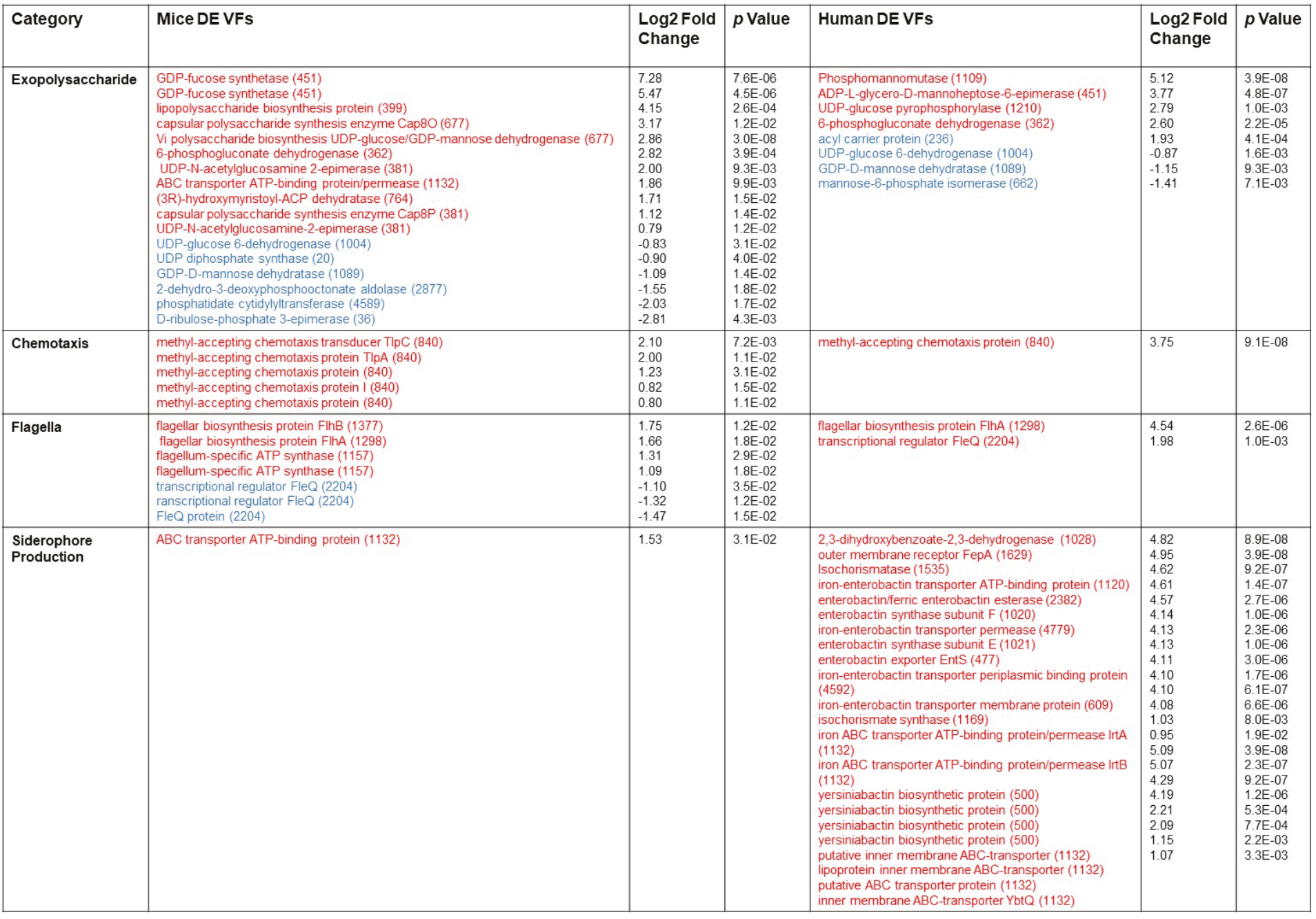
Specific aging-associated differential virulence factors in the murine and human gut microbiome. VFDB hits from Fig 3 and Fig 5 are listed by species (mouse vs human), virulence factor category, and association with aging (red = overabundant in aged, blue = overabundant in young). Log-fold change and *p* value are listed for each individual gene.

**Data file S1.** Virulence Factor Database annotation of complete blood resistant *E. coli* (JC001) and *E. faecalis* JC002) genomes including COG number.

**Data file S2.** Comparison clinical *E. coli* and *E. faecalis* isolates utilized for comparative genomic and phylogenetic tree analyses.

